# *Schistosoma mansoni* infection enhances insertion sequence activity in the host gut microbiome

**DOI:** 10.64898/2026.09.23.753824

**Authors:** Yi-Hsuan Lee, Gabriel Rinaldi, Alba Cortes, Cinzia Cantacessi, Wayne Aubrey, Martin T. Swain

## Abstract

*Schistosoma mansoni* infection is associated with alterations of the host gut microbial community structure, yet its impact on microbial genome dynamics and horizontal gene transfer remains poorly understood. In this study, we investigated the activity of Mobile Genetic Elements (MGEs), and of Insertion Sequences (IS) in particular, in the gut microbiome of wild-type (WT) and human-microbiota-associated (HMA) mice. Taxonomic profiling revealed significant differences in bacterial diversity between WT and HMA mice, with reduced richness and alpha diversity observed in HMA gut microbiomes regardless of the infection status. Principal Coordinates Analysis demonstrated clear separation between microbial communities, reflecting differences in host background and bacterial composition. In contrast, differences were observed between plasmid and viral community profiles and between these and bacterial diversity, suggesting that different MGEs may follow distinct ecological dynamics within the gut microbiome. IS elements within bacterial genomes were widespread across all microbiome communities analysed in this study, with higher insertion counts observed not only in HMA compared to WT mice, but also in infected compared to uninfected mice. IS elements were predominantly associated with dominant gut bacterial taxa, particularly members of *Bacteroidia* and *Clostridia*. At the genomic level, insertion events occurred more frequently in intergenic regions than in coding sequences, suggesting a preferential selective pressure towards alteration of gene expression rather than gene disruption. Furthermore, IS activity was dominated by a subset of IS families, with consistent patterns observed across experimental groups. Notably, infection was associated with increased insertion frequencies across multiple taxa and IS families, suggesting enhanced mobilisation under conditions of host-associated stress (the physiological and molecular pressure experienced by microorganisms, pathogens, or the host itself during their interactions within a symbiotic or infected environment) and microbiome perturbation. Together, these findings demonstrate that *S. mansoni* infection may promote IS mobilisation in the mouse gut microbiome, potentially contributing to microbial genome plasticity and facilitating horizontal gene transfer within host-associated microbial communities.

## Introduction

The gut microbiome is a complex and dynamic ecosystem that plays a central role in homeostasis, metabolism, and immune regulation. Within this ecosystem, microbial communities are continuously shaped by host factors, environmental conditions, and interactions among microbial taxa ^1^. One key mechanism underlying microbial interaction is horizontal gene transfer (HGT), which enables the rapid acquisition and dissemination of genetic material across microbial populations ^1,2^. Mobile genetic elements (MGEs), including plasmids, bacteriophages, and insertion sequences (IS), are major players in this process ^3,4^.

IS are among the simplest MGEs, typically encoding only the machinery required for transposition ^4^. Despite their minimal structure, IS elements can substantially impact microbial genomes by mediating gene disruption, promoting genomic rearrangements, and modulating gene expression through insertion into regulatory regions ^5,6^. The activity of IS elements is often influenced by environmental stress, suggesting that they may play an important role in microbial adaptation under changing ecological conditions ^5,7^. Given that IS elements can facilitate genome plasticity and contribute to HGT, changes in their activity may impact microbial adaptation, community stability, and the spread of functional traits such as antimicrobial resistance (AMR) ^4^. This is particularly relevant during inflammatory processes, such as those associated with infection.

Helminth infections represent a major source of perturbation within the gut ecosystem ^8,9^. Infection with the blood fluke *Schistosoma mansoni*, a causative agent of hepato-intestinal schistosomiasis that affects more than 250 million people globally ^10^, has been associated with significant alterations in the gut microbiome composition and/or functional capacity in naturally parasitised humans and rodent models of infection ^11–13^. Previous work using human-microbiota-associated (HMA) mice demonstrated that human-derived microbial communities exhibit altered colonisation dynamics, leading to reduced bacterial diversity in the mouse gastrointestinal tract upon *Schistosoma* infection. Moreover, substantial differences in infection-associated changes in gut microbiota composition were observed in HMA compared to wild type (WT) mice ^14–16^. Whilst taxonomic and functional changes in bacterial communities during infection have been increasingly well characterised, the impact of the infection on other (non-prokaryotic) members of the gut flora, as well as on microbial genome dynamics and MGE activity, remains poorly explored ^3,5,17^.

In this study, we analysed shotgun metagenomic data from WT and HMA mice, in the presence or absence of *S. mansoni* infection aiming to: (1) characterise the impact of IS across gut microbial taxa, (2) assess differences in insertion activity between host basal microbiome backgrounds and infection status, (3) map the insertion site affinity within bacterial genomes, and (4) study the impact of the infection in other organisms associated with the host gut, including protozoa, fungi, and viruses. By integrating taxonomy, genomics and analysing IS activity, this work provides new insight into how host infection and microbiome ecosystem shape MGE activity in the gut.

## Results

### Gut microbial profiles vary between WT and HMA mice

Whole genome sequencing (WGS) data generated in our previous study ^15^ were obtained and analysed in the current study; briefly, WGS was performed from DNA extracts from intestinal content of *S. mansoni*-infected WT mice (hereafter referred to as WP for ‘Wild Positive’), *S. mansoni*-uninfected WT mice (WN for ‘Wild Negative’), *S. mansoni*-infected HMA mice (HP for ‘Humanised Positive’), and *S. mansoni*-uninfected HMA mice (HN for ‘Humanised Negative’) ^15^. Taxonomic profiling of the filtered metagenomic data in the present analysis detected a total of 7,567 bacterial features (i.e., taxonomically distinct bacterial taxa identified by the profiling Kraken2, Bracken, and MicrobiomeAnalyst pipelines) across all samples, encompassing both WT (7,513 species) and HMA mice (7,117 species) (Supplementary Table S1). In addition to bacteria, sequences assigned to plasmids (features: 966; WT: 918 species; HMA: 828 species) (Supplementary Table S2), viruses (features: 109; WT: 92 species; HMA: 87 species) (Supplementary Table S3), fungi (features: 80; WT: 50 species; HMA: 50 species) (Supplementary Table S4), and protozoa (features: 36; WT: 16 species; HMA: 16 species) (Supplementary Table S5) were also identified (Supplementary Table S6).

Bacterial species richness and alpha diversity, measured by the Shannon index, were significantly higher in WT mice than in HMA mice (Fig. S1A) and differed significantly across experimental groups, albeit not within the same mouse line (HMA or WT mice) (Fig. S1B). A marked decrease in measured gut bacterial species richness was previously observed in HP mice ^15,18^ (Fig. S1C). These findings are consistent with previous studies employing both shotgun metagenomics (species richness) and 16S rRNA sequencing (Shannon index) approaches ^15,19^.

Principal Coordinates Analysis (PCoA) revealed significant separation between HMA and WT gut bacterial communities, irrespective of infection status (Fig. S1D). This separation likely reflects differences not only between the original gut microbiota inoculated in the parental generation of HMA mice compared to WT, but also differences in microbiome establishment following human microbiota transplantation, as well as breeding and housing conditions between the two mouse lines (i.e., maintenance of WT mice under specific pathogen-free conditions versus isolation of HMA mice) ^20^.

Plasmids and viruses coexist with bacterial genomes within the cell; however, species richness and alpha diversity for plasmids and viruses showed trends that differed from those observed for bacterial taxa. Plasmid species richness and alpha diversity differed significantly across experimental groups, with HP mice showing significantly higher values than HN mice (Fig. S2A). In addition, PCoA revealed clear separation between HMA and WT gut plasmid communities, irrespective of infection status (Fig. S2B).

Viral taxa richness and alpha diversity also differed significantly across experimental groups, with HN mice showing significantly lower values than the other groups (Fig. S2C). Likewise, PCoA showed clear separation between HMA and WT gut viral communities, irrespective of infection status (Fig. S2D).

Fungal species richness and alpha diversity showed a similar trend to that observed for bacterial communities. WT mice exhibited significantly higher fungal richness and alpha diversity than HMA mice, and values differed significantly across experimental groups (Fig. S3A). This pattern is consistent with a reduced capacity of selected human-associated microbes to colonise the mouse gastrointestinal tract. PCoA showed partial separation between HMA and WT gut fungal communities, irrespective of infection status (Fig. S3B).

In contrast, protozoan species richness and alpha diversity did not differ significantly across experimental groups (Fig. S3C). PCoA showed limited separation between HMA and WT gut protozoan communities (Fig. S3D).

Taken together, these results indicate that bacterial communities were strongly affected by both microbiome humanisation and infection status, whereas plasmids and viruses exhibit distinct and partially decoupled dynamics, with the protozoan component of the gut microbiome remaining comparatively stable. This domain-specific response highlights that changes in bacterial community structure do not necessarily result in parallel shifts in the associated plasmid, viral, and eukaryotic communities.

### IS are widespread in the gut microbiota of both WT and HMA mice

IS elements were identified in the gut metagenomes of *S. mansoni*-infected and -uninfected WT and HMA mice using the pseudoR pipeline ^21^. All groups showed evidence of substantial heterogeneity in IS. In total, 1,149 IS were detected in WN mice, 1,775 in WP mice, 675 in HN mice, and 780 in HP mice (Fig. 1A). Among these, 648 IS in open reading frames (ORFs) were detected in WN mice, 1,005 in WP mice, 337 in HN mice, and 445 in HP mice (Fig. 1B).

**Fig. 1:**
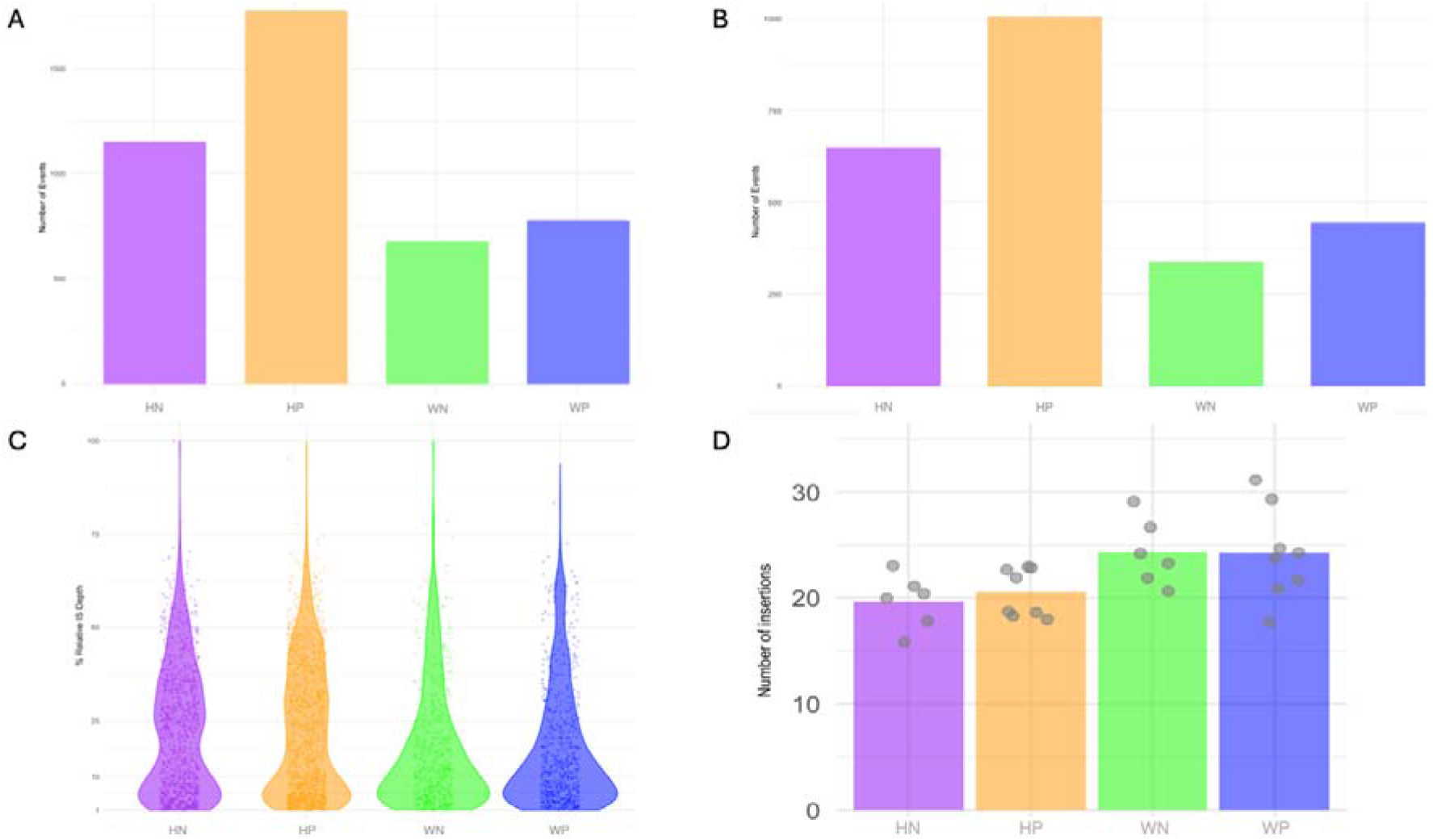
IS are widespread in the gut microbiome of S. mansoni-infected and -uninfected WT and HMA mice microbiota. (A) Sum of number of insertions per sample. (B) Sum of insertions in ORF per sample. (C) Violin plot for abundance of IS in the gut microbiome of S. mansoni-infected and -uninfected WT and HMA mice. Each data point represents one new insertion not found in the reference assembly. The y-axis is the relative IS depth percentage. Each violin represents the kernel density of IS abundance values within a group, with wider sections indicating higher data density, and overlaid points represent individual observations. (D) Number of unique IS per sample. Each dot represents individual mice within the groups.

Relative IS depth was used to quantify the abundance of IS insertion alleles ^21^. Across all groups, relative IS depth values were positively skewed, with a high concentration of low-abundance IS and a smaller number of high-abundance outliers (Fig. 1C). HN and HP showed broader distributions with higher densities in the mid-abundance range, whereas WN exhibited a stronger concentration of low-abundance IS and fewer high-abundance observations. WP showed an intermediate distribution with moderate spread and several high-abundance IS (Fig. 1C).

### *Schistosoma mansoni* infection is associated with increased IS activity

Remarkably, between both mouse lines, infected groups exhibited higher numbers of IS than their uninfected counterparts. In addition, HMA mice showed higher IS counts than WT mice (Fig. 1C). We next quantified unique IS per sample within each group. The mean relative abundance of unique IS differed among experimental groups, with HN, HP, WN, and WP showing distinct insertion frequencies (Fig. 1D). Taken together, this pattern suggests an association between an increased mobilisation of IS in the gut microbiome and murine schistosomiasis. Altered microbiome composition may be affected by a reduced colonisation capacity of human-associated microbes in the mouse gastrointestinal tract, as well as environmental stress and/or community-level instability within the gut microbiome during schistosomiasis ^14,15^.

IS activity was unevenly distributed across bacterial taxa in the gut microbiome. At the class level, the majority of IS were associated with members of *Bacteroidia* and *Clostridia*, which showed the highest insertion frequencies across all experimental groups. Additional insertional activity was observed in classes such as *Bacilli*, *Gammaproteobacteria*, and *Betaproteobacteria*, although at lower frequencies. In contrast, classes including *Cytophagia*, *Deferribacteres*, *Epsilonproteobacteria*, *Erysipelotrichia*, *Flavobacteriia*, and *Tissierellia* showed relatively few insertion events (Fig. 2; Fig. S4).

**Fig. 2:**
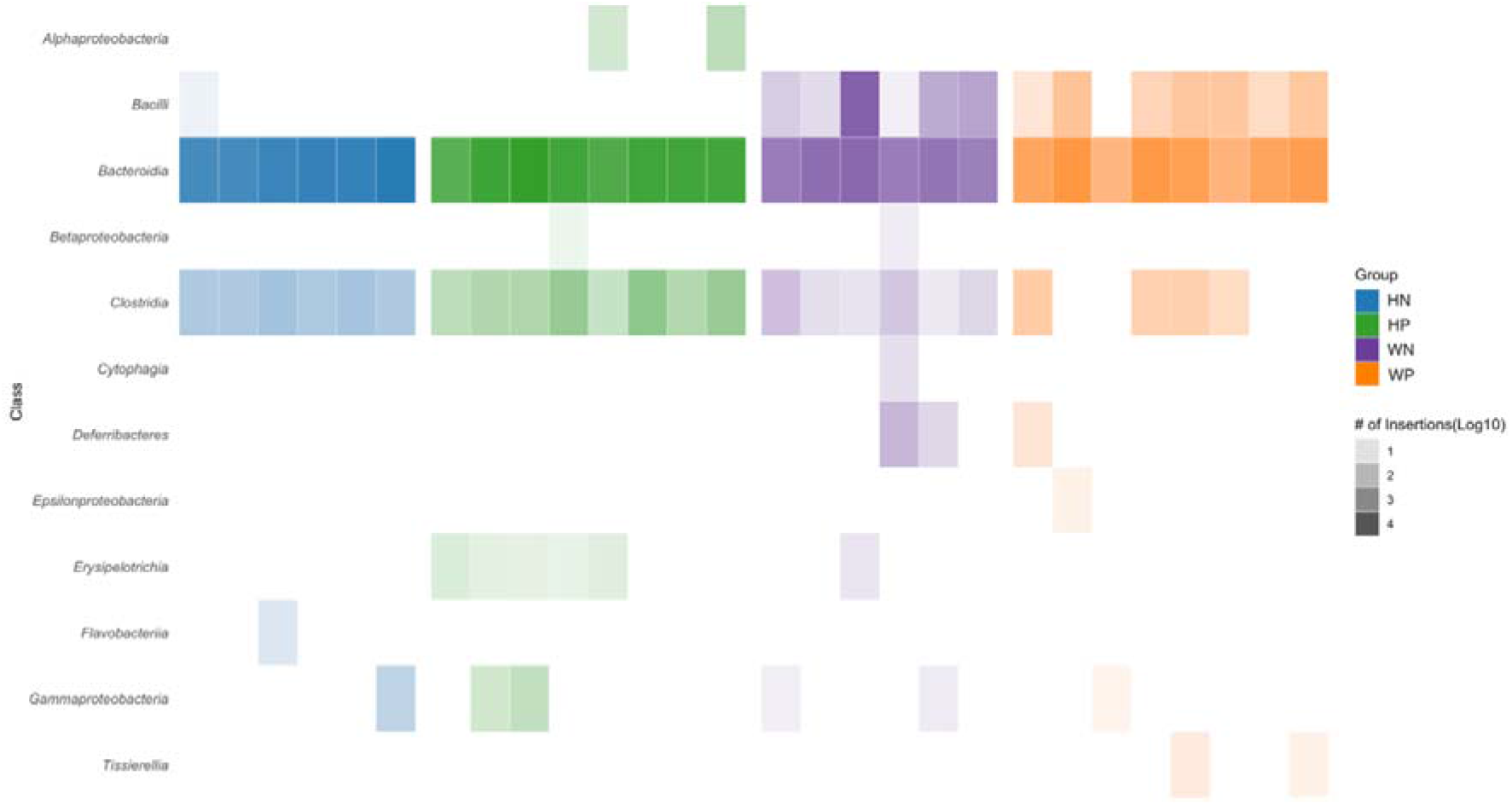
IS per bacterial class. Each column represents data collected from individual mice across the four experimental groups as indicated. The intensity of each bar is proportional to the number of IS as indicated.

At genus level, IS were concentrated within several dominant gut taxa. Genera such as *Bacteroides*, *Phocaeicola*, *Parabacteroides*, *Prevotella*, *Blautia*, and *Ruminococcus* accounted for a large proportion of insertion events across samples (Fig. S4). These genera are major constituents of the mammalian gut microbiota and therefore provide extensive genomic landscape targets for IS mobilisation. Additional insertions were detected in genera including *Odoribacter*, *Muribaculum*, *Coprobacter*, and *Alistipes*, although these occurred at lower frequencies (Fig. S4).

Clear differences in IS patterns were also observed amongst experimental groups; HP mice showed a broader distribution and higher abundance of IS across multiple taxa compared to HN mice, particularly within *Bacteroidia*-associated genera such as *Bacteroides*, *Phocaeicola*, and *Parabacteroides* (Fig. S4). A similar trend was observed in the WT mouse line, where WP mice exhibited more widespread insertion events than WN mice. In both mouse lines, infection was therefore associated with increased IS activity across dominant gut bacterial taxa.

Comparisons between mouse lines also revealed differences in the taxa affected by IS. HMA mice (both HN and HP) showed relatively concentrated insertion activity within a smaller number of dominant genera, whereas WT mice (both WN and WP) exhibited insertions distributed across a broader range of taxa, including several lower-abundance classes (Fig. 2; Fig. S4). Nevertheless, across all four groups, the overall pattern remained consistent, with the majority of IS activity occurring within *Bacteroidia* and *Clostridia* lineages (Fig. 2; Fig. S4). Notably, *Erysipelotrichia* showed IS activity exclusively in infected HMA mice, with insertions detected in 5 of 8 infected mice compared with none of the 6 uninfected HMA controls. This group-specific pattern suggests that IS activity within *Erysipelotrichia* may be associated with the altered gut environment during infection (Fig. 2; Fig. S4).

Together, these results indicate that IS mobilisation may be strongly associated with dominant gut bacterial taxa and that both infection status and host microbiome background influence the distribution and frequency of insertion events within the gut microbial community.

Although IS within *Bacteroidia* were detected across all experimental groups, the specific IS elements contributing to these insertions varied substantially between groups (Fig. 3A; Fig. S4; Fig. S5A). Several IS families, including IS4, IS5, IS66, IS982, and IS21, were frequently associated with taxa within the *Bacteroidia*; however, individual IS elements within these families displayed distinct group-specific patterns. Remarkably, multiple IS4- and IS66-associated elements showed strong insertion signals in the infected groups (HP and WP), compared to uninfected control mice (Fig. 3; Fig. S4 and S5). These observations suggest that while the broader IS family repertoire is shared across host conditions, infection status may influence the activity or expansion of specific IS elements within dominant *Bacteroidia* lineages.

**Fig. 3:**
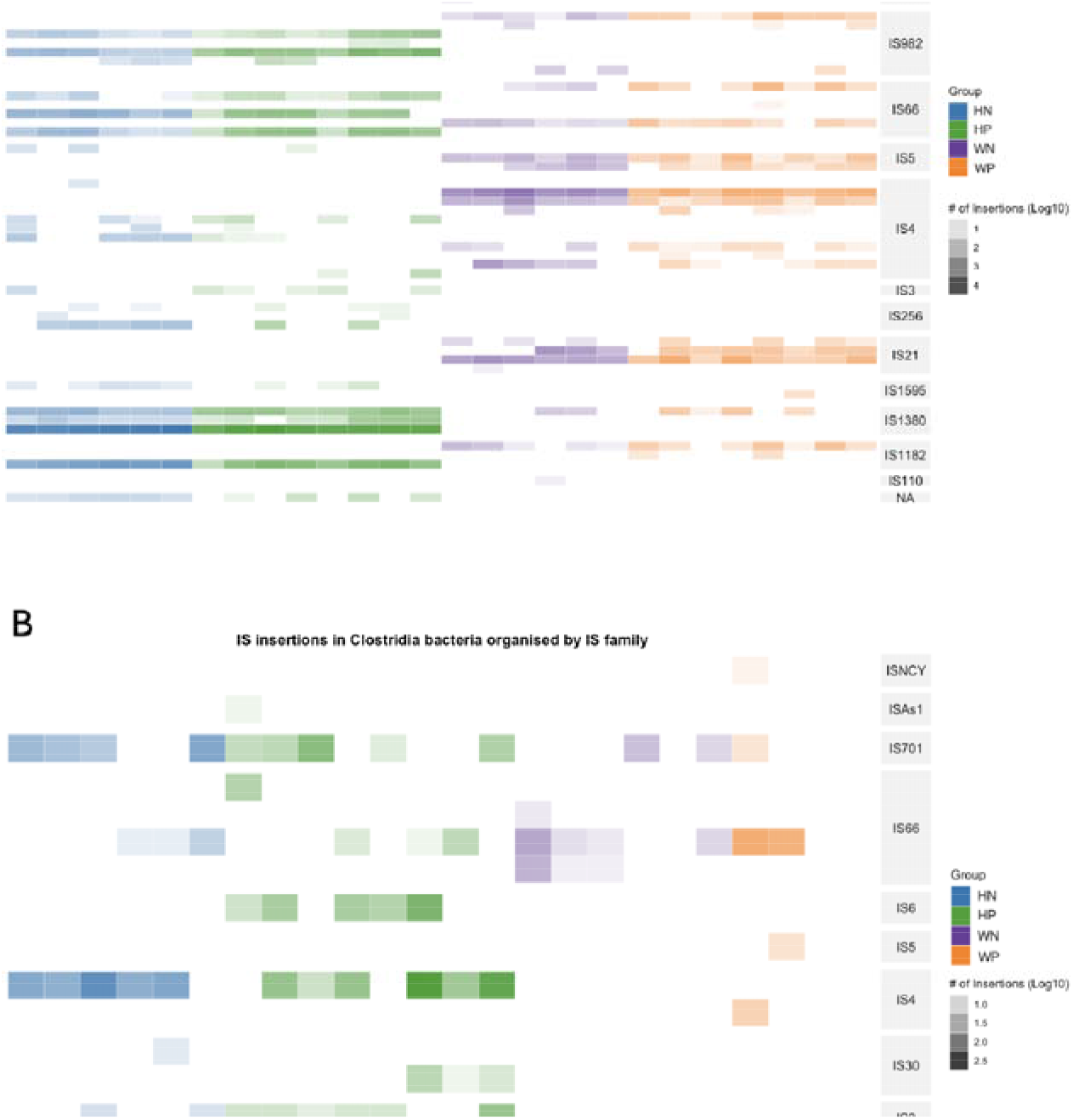
IS in Bacteroidia and Clostridia bacteria organised by IS family. The intensity of each bar is proportional to the number of IS.

A similar pattern was observed for *Clostridia*, where IS were also detected across all groups but were dominated by a smaller subset of IS families, including IS4, IS66, IS701, and IS1595 (Fig. 3B; Fig. S4 and S5B). In particular, IS4-related elements showed frequent insertions across both HMA and WT mouse lines, irrespective of the infection status, while IS66 and IS1182 insertions were more abundant in infected samples. Notably, several IS families were shared between *Bacteroidia* and *Clostridia*, including IS4 and IS66, although the specific IS elements differed between the two bacterial classes. This may suggest that while certain IS families may be broadly distributed and capable of mobilising across multiple bacterial taxa, the individual IS elements themselves appear to be lineage-specific.

In summary, these results suggest that IS activity in the gut microbiome is structured both by bacterial phylogeny, baseline gut microbiota and infection status. Dominant gut bacterial classes such as *Bacteroidia* and *Clostridia* harbour diverse but partially overlapping repertoires of IS families, with infection-associated microbiome changes potentially promoting the expansion or mobilisation of selected IS elements.

IS elements can insert into either coding sequences (intragenic regions) or intergenic regions of bacterial genomes, where they may disrupt gene function or alter the expression of adjacent genes, respectively ^5,6,22^. To compare insertion frequencies across different genomic regions and experimental groups while accounting for differences in the amount of genomic sequence available for insertion, we calculated the occurrence per million base pairs (OPM) normalised insertion rate (see Methods).

Intragenic and intergenic insertion frequencies were compared across the four experimental groups (HN, HP, WN, and WP) and different IS families (Fig. 4). Across all groups, intergenic insertions were generally more frequent than intragenic insertions for most IS families. This pattern was particularly evident for IS1380, IS4, ISL3, and IS3, which showed consistently higher numbers of intergenic insertions. Several additional elements, including IS21, IS66, and ISAs1, also contributed substantially to the overall insertion counts (Fig. 4).

**Fig. 4:**
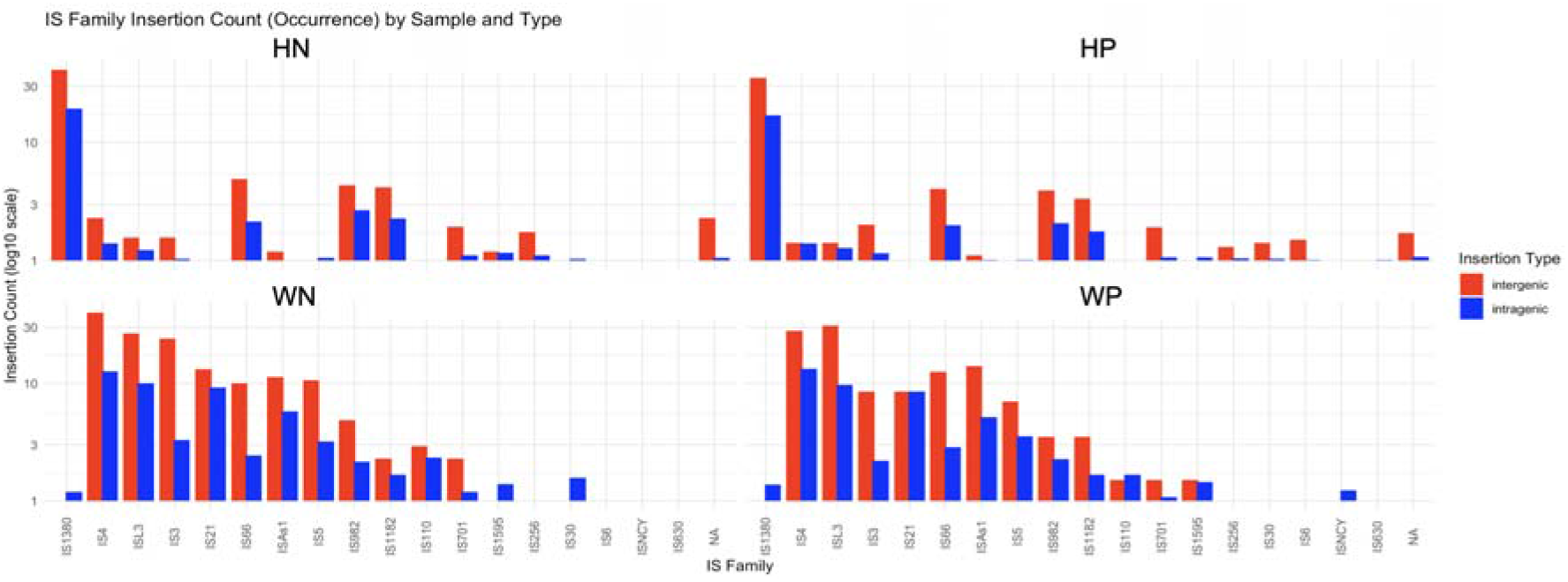
Preferential IS in intergenic (red) and intragenic (blue) regions, shown as IS family occurrences per million base pair loci.

Differences in insertion patterns were also observed between experimental groups. In HMA mice, infected animals (HP) showed slightly higher insertion rates across several IS families compared to uninfected animals (HN) (Fig. 4). A similar trend was observed in WT mice, where infected mice (WP) exhibited higher insertion rates than uninfected mice (WN) (Fig. 4). However, WT mice displayed a broader distribution of insertion events across multiple IS families, particularly IS4, ISL3, IS3, and IS21, whereas insertions in HMA mice were more concentrated within a smaller subset of families (Fig. 4).

These results indicate that IS activity in the mouse gut microbiome is dominated by a limited number of IS families, which more frequently occurred in intergenic regions than within coding sequences. This observation suggests that many IS elements preferentially integrate into genomic regions in which protein coding sequences are not disrupted. Given that intragenic IS can disrupt gene function, the lower frequency of such insertions may partly reflect purifying selection against deleterious insertion events, although the present analysis cannot distinguish this process from intrinsic insertion-site preferences ^23,24^. On the other hand, IS in intergenic regions may also influence the expression of neighbouring genes, for example through effects on local regulatory architecture and promoter activity ^25,26^.

### IS preferentially target adaptive functional categories

Having identified extensive IS across predicted coding sequences, we next characterised ORFs containing insertions (iORFs) across the four sample groups (HN, HP, WN, WP). A deduplicated ORF catalogue was used to distinguish between shared iORFs (present across multiple samples) and unique iORFs (restricted to individual samples). Across all groups, shared iORFs were substantially more abundant than unique iORFs, with HP showing the highest number of shared insertions, followed by HN, WP, and WN. In contrast, unique iORFs were consistently low across all groups. This pattern suggests that a core set of insertion targets is conserved. However, a small subset of IS may have also contributed to sample-specific variation (Fig. 5).

**Fig. 5:**
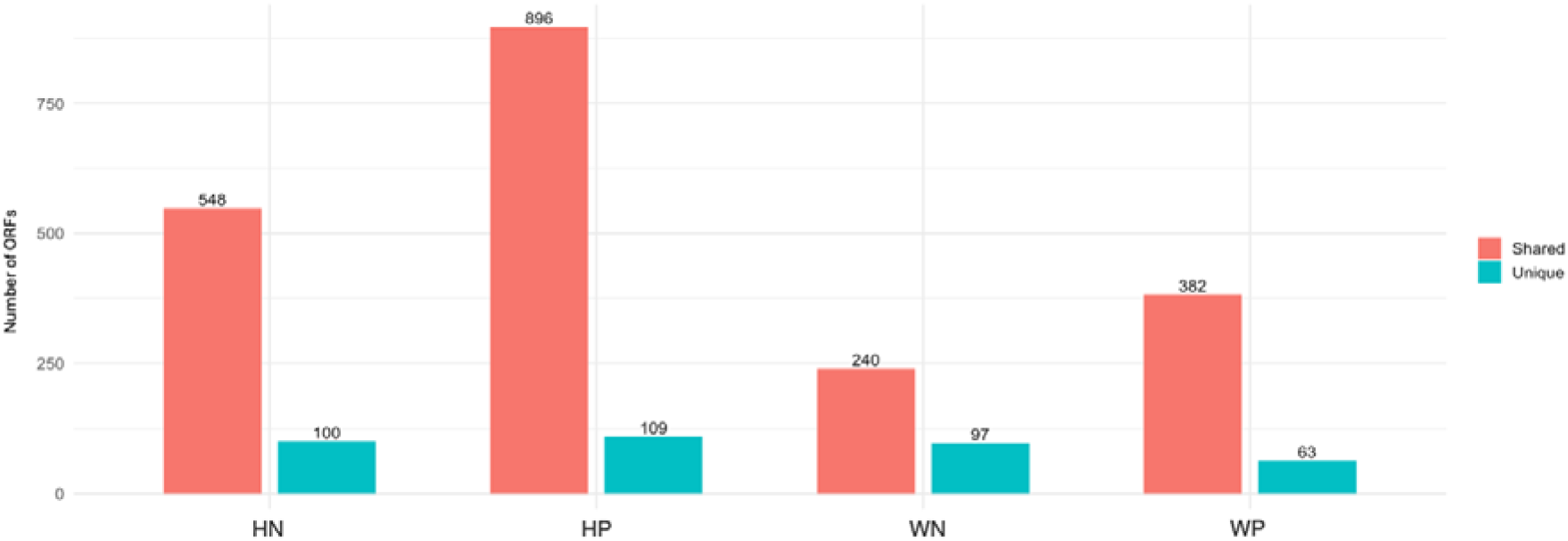
Number of shared and unique open reading frames containing insertions (iORFs) within HP, HN, WP, and WN individuals

To further explore potential functional perturbations associated with the characterised IS, iORFs were grouped into broad functional categories (Fig. 6). Insertions were most frequently observed in genes associated with substrate uptake systems, particularly susC/susD and TonB-dependent receptors, which mediate the acquisition of nutrients and other substrates and may contribute to metabolic flexibility under changing environmental conditions. These systems are closely linked to the bacterial cell envelope, with SusC functioning as an outer-membrane transporter and SusD as an associated substrate-binding lipoprotein, while TonB-dependent receptors facilitate the uptake of scarce nutrients across the outer membrane (REFS). Consistent with this pattern, genes involved in cell wall, membrane, and envelope biogenesis also showed substantial insertion frequencies, suggesting that membrane-associated functions represent important targets of MGE insertion and may influence bacterial adaptation to environmental and host-associated conditions (Fig. 6). Anti-MGE defence systems exhibited moderate insertion levels, indicating interactions between MGEs and host defence mechanisms that may shape genome stability and the persistence of mobile elements. AMR-associated genes showed comparatively lower but consistent insertion frequencies across samples (Fig. 6), suggesting that although resistance determinants were less frequently targeted, their association with mobile genomic regions may facilitate their persistence and potential dissemination within bacterial communities ^26,27^.

**Fig. 6:**
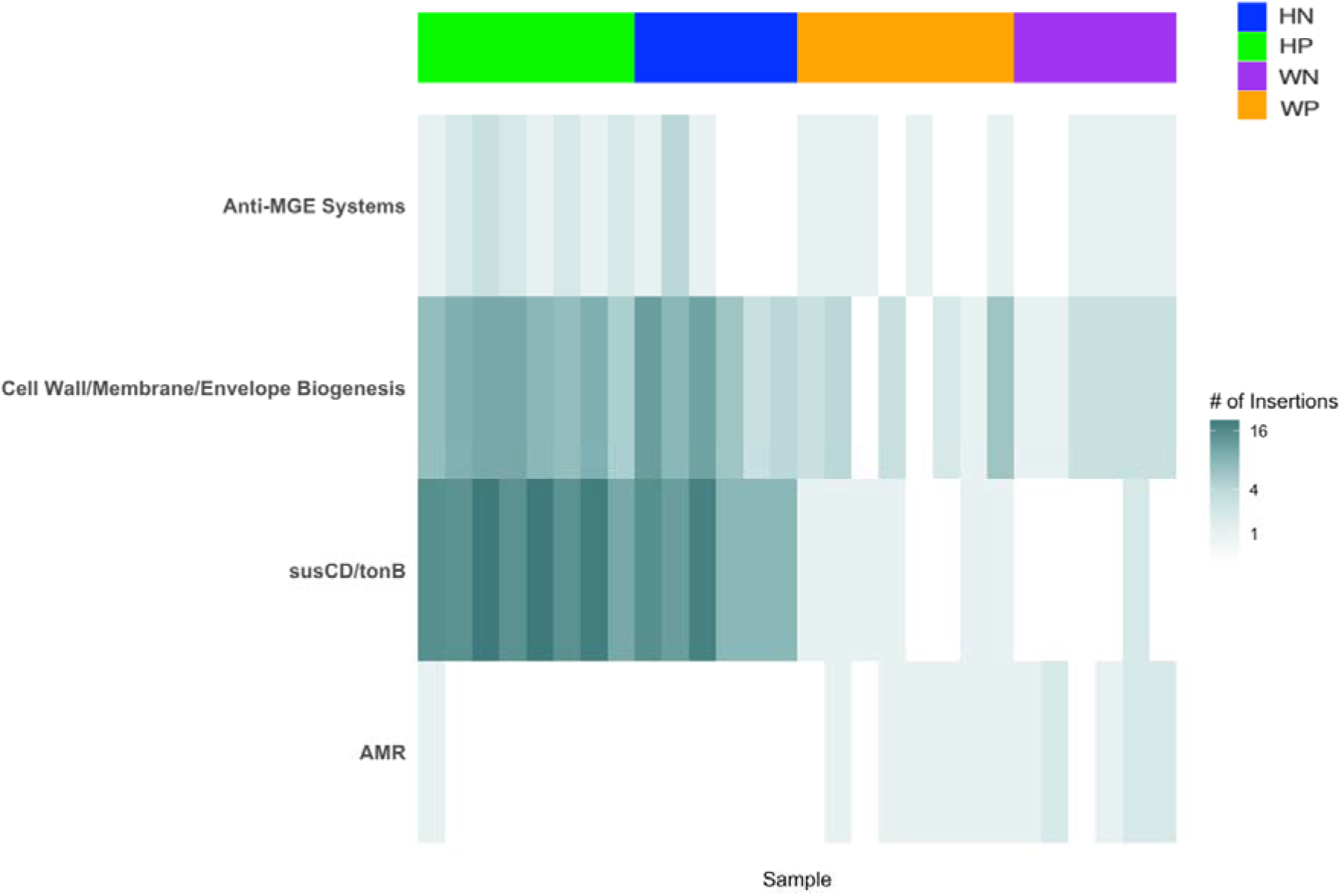
Heatmap of IS in four discrete gene functional categories as indicated across the four experimental groups.

Notably, the distribution of insertions varied between sample groups, with HP and HN generally exhibiting higher insertion densities across most functional categories compared to WN and WP (Fig. 6). This suggests that infection status and/or host condition may influence both the frequency and functional targeting of IS elements.

Overall, these results demonstrate that IS elements preferentially insert into genes associated with nutrient acquisition, cell surface modification, and genomic defence, supporting their role in shaping bacterial adaptability. At the same time, the coexistence of shared and unique iORFs highlights a balance between conserved insertion hotspots and individual-specific genomic variation within the gut microbiome.

## Discussion

In the present report, we studied microbiome data from WT and HMA mice in the presence or absence of *S. mansoni* infection. The observed differences in bacterial alpha diversity were consistent with previous studies, which reported significantly higher diversity in WT compared with HMA mice, tentatively attributed to a reduced capacity of selected human gut microbes to colonise the mouse gastrointestinal tract ^14,15^. In addition to colonisation constraints, the present metagenomic analysis suggests environmental stress, such as that associated with parasite infections, or community-level instability within the HMA gut microbiome, may also contribute to the reduced bacterial diversity observed. It is also important to consider methodological differences when interpreting diversity estimates between studies. The Shannon index derived from 16S rRNA gene amplicon sequencing may differ from that obtained through shotgun metagenomics due to several factors, including primer bias that affects amplification efficiency, differences in taxonomic resolution between methods, variation in 16S rRNA gene copy numbers across bacterial taxa, and differences in bioinformatic pipelines, reference databases, and filtering strategies ^28^. Furthermore, sequencing depth can influence the detection of rare taxa, thereby affecting diversity estimates. These methodological considerations may at least partially account for quantitative differences between 16S-based and shotgun metagenomic analyses, while the overall trends remain biologically consistent across approaches ^28^. Notably, the distinct responses observed for plasmids and viruses indicate that changes in bacterial diversity do not necessarily translate directly into parallel changes in associated MGEs and bacteriophages. The observed differences in bacterial alpha diversity are therefore unlikely to fully explain the patterns detected for these elements. Instead, the dynamics of MGEs and bacteriophages may additionally reflect altered rates of HGT, phage induction, and stress-associated mobilisation within the gut microbiome ^3,17,29,30^. This partial decoupling is particularly relevant to the present study because environmental perturbation associated with *S. mansoni* infection may influence not only the composition of bacterial communities but also the activity of MGEs independently of changes in bacterial richness or alpha diversity. Further studies are needed to fully understand the biological implications of these findings.

To investigate how microbiome humanisation coupled with experimental *S. mansoni* infection may influence bacterial genome plasticity and HGT dynamics, we analysed the IS activity across the gut microbiome using metagenomic sequencing data. IS elements are among the most abundant MGEs in bacterial genomes and can drive genomic rearrangements, gene disruption, and the mobilisation of genetic material between microbial populations. IS activity in the gut microbiome showed clear taxonomic, genomic, and host-associated patterns, providing insight into how *S. mansoni* infection and microbiome humanisation influence HGT dynamics. Across all samples, IS were significantly concentrated within dominant gut bacterial lineages, particularly members of *Bacteroidia* and *Clostridia*. These taxa represent major components of the intestinal microbiota and therefore provide extensive genomic substrates for IS mobilisation. The enrichment of IS within these groups suggests that insertional activity is strongly shaped by microbial abundance and ecological dominance, rather than random distribution across taxa ^3,5^. The detection of IS within *Erysipelotrichia* exclusively in infected HMA mice is also noteworthy. Although the present study cannot establish a causal relationship between *S. mansoni* infection and IS activity in this taxon, the infection-specific detection may indicate that changes in the intestinal environment during murine schistosomiasis create ecological conditions under which IS activity in members of *Erysipelotrichia* becomes more apparent. Members of *Erysipelotrichia* have been associated with host-microbiome interactions and inflammatory processes, providing a potential biological context for the infection-specific IS activity observed in this study ^31^. However, whether increased IS activity contributes to altered functions within *Erysipelotrichia* or instead reflects changes in the abundance, composition, or ecological conditions affecting these bacteria remains unresolved. Further studies combining longitudinal sampling, measurements of bacterial abundance, and functional characterisation of IS-associated genes will be required to determine whether IS activity in *Erysipelotrichia* contributes to bacterial adaptation during schistosome infection.

At the genomic level, observed IS elements were more frequently associated with intergenic regions rather than with coding sequences. This pattern was consistent across IS families and experimental groups and may reflect either preferential insertion into non-coding regions, selective removal of deleterious insertions in coding sequences, or a combination of both processes. Because the present analysis captures surviving insertion events rather than the initial transposition events, it is not possible to distinguish between intrinsic insertion-site preferences and subsequent selection against deleterious insertions ^32^. Nevertheless, intergenic IS may influence the expression of neighbouring genes while reducing the likelihood of directly disrupting protein-coding sequences. Such regulatory effects could contribute to microbial adaptation and genome plasticity within the gut environment ^21^. In contrast, coding sequence insertions, although less frequent, are more likely to result in gene disruption, altered protein function, or other phenotypic consequences that may be subject to stronger selective pressures ^5,6,33^.

IS integration patterns varied across IS families, indicating that insertional behaviour is, at least in part, element-specific ^5,6,34^. A subset of IS families, including IS1380, IS4, ISL3, and IS3, accounted for a large proportion of insertion events, suggesting that these elements may have greater transpositional activity, persistence, or ecological fitness within the gut environment. Several of these families have also been associated with functional consequences beyond genome rearrangement, including disruption or modulation of neighbouring genes and, for some members, the mobilisation or altered expression of antimicrobial resistance determinants. In particular, IS1380, IS3, and IS4 family members have been reported to influence the expression or mobilisation of adjacent resistance genes, whereas ISL3 elements have been implicated in genomic rearrangements and insertion-associated changes in gene structure. The enrichment of these IS families in the present dataset may therefore reflect not only differences in transposition mechanisms and target-site preferences but also their capacity to interact with functionally important regions of bacterial genomes ^4,35–39^. The presence of both broadly distributed and low abundance IS families further supports a model in which different elements contribute unequally to genome diversification, potentially reflecting differences in transposition mechanisms, target site preferences, or regulatory control ^5,6,33^.

Importantly, both infection status and host microbiome background influenced IS dynamics. Infected mice consistently exhibited higher insertion counts than their uninfected counterparts within both HMA and WT lines, indicating that *S. mansoni* infection might be associated with increased IS mobilisation. This may reflect infection-induced alterations in the gut environment, including inflammation, immune activation, or altered nutrient availability, which could contribute to changes in bacterial community structure and create conditions favourable for IS mobilisation ^5,7,9,13,15^.

Stress conditions are known to activate transposition and other MGE, thereby increasing genome plasticity and the potential for rapid adaptation ^36^.

Differences between HMA and WT mice further suggest that microbiome composition and stability play a key role in shaping IS activity. HMA mice, which harbour human-derived microbial communities, showed patterns consistent with increased insertional activity concentrated within a smaller subset of taxa. This may reflect reduced colonisation efficiency or ecological instability in the mouse gut, leading to heightened stress and increased mobilisation of IS elements ^5,7,14,40^. In contrast, WT mice exhibited a broader distribution of IS across taxa, consistent with a more stable and diverse microbial community in which insertional events are more widely dispersed.

The combined effects of infection and microbiome context suggest that IS mobilisation is driven by both ecological disturbance and host-associated factors ^3,5,9,13^. Increased insertion activity in infected and humanised microbiomes supports the hypothesis that IS elements contribute to microbial adaptation under conditions of environmental stress and community restructuring. These dynamics are likely to have downstream consequences for HGT, including the dissemination of accessory genes such as AMR determinants and metabolic functions ^21^. Functional annotation of iORFs further suggests that IS activity involved genes associated with nutrient acquisition, microbial competition, host interaction, and genome defence. Because these analyses were restricted to iORFs, the identified functional categories represent coding sequences directly disrupted by IS rather than genes that may be affected indirectly through changes in neighbouring gene regulation. In particular, iORFs associated with SusCD/TonB transport systems, exopolysaccharide biosynthesis, and anti-mobile genetic element (Anti-MGE) defence mechanisms suggest that IS may directly affect functions involved in nutrient acquisition, microbial fitness, and defence against mobile genetic elements. Consequently, IS mobilisation within these coding sequences may contribute to changes in bacterial fitness and ecological interactions within the gut environment, although the functional consequences of individual insertions require experimental validation.

Beyond their potential effects on overall genome plasticity, the functional distribution of iORFs provides further insight into the potential consequences of IS activity. The preferential occurrence of IS in genes involved in substrate uptake, cell envelope biogenesis, Anti-MGE defence, and AMR suggests that mobile elements preferentially target gene functions involving interactions between bacterial cells and their surrounding environment ^21^. In particular, the enrichment of insertions in susC/susD and TonB-dependent receptor systems is notable because these systems mediate the acquisition of nutrients and other substrates at the bacterial cell envelope and can contribute to metabolic flexibility under changing environmental conditions. SusC functions as an outer-membrane transporter, whereas SusD is an associated substrate-binding lipoprotein, while TonB-dependent receptors facilitate the uptake of specific nutrients and other scarce substrates across the outer membrane. Alteration of these loci could therefore influence substrate utilisation and competitive interactions within the gut microbiome. Similarly, the relatively high frequency of insertions in genes involved in cell wall, membrane, and envelope biogenesis may indicate that surface-associated functions are important targets or potential consequences of IS activity, with possible implications for envelope integrity and adaptation to host-associated conditions. The detection of insertions within Anti-MGE defence systems further highlights a potential evolutionary interaction between mobile elements and bacterial mechanisms that restrict their propagation. Consistent with this interpretation, IS transposition has been shown to disrupt CRISPR-Cas immunity through insertion into Cas genes and to target other bacterial defence systems ^41^. Although AMR-associated genes showed lower insertion frequencies, their consistent detection indicates that resistance-associated loci remain connected to the broader mobile genetic landscape and may represent potential targets for future investigation of IS-mediated mobilisation. Together, these observations suggest that IS elements are not distributed randomly across bacterial genomes but may disproportionately affect genomic regions involved in nutrient acquisition, cell-envelope functions, environmental interactions, and defence against mobile DNA. Such insertional activity could contribute to bacterial adaptation and genome plasticity within the gut environment ^21,26,41^.

In addition to bacteria, this study characterised plasmid, viral, fungal, and protozoan components of the gut metagenome. While bacterial responses to schistosomiasis have been examined previously, little is known about how these other microbial domains respond to infection or microbiome humanisation. Plasmid and viral communities displayed patterns distinct from those observed for bacterial taxa, suggesting that the dynamics of mobile genetic elements and bacteriophages are not solely determined by bacterial diversity. In contrast, fungal communities broadly mirrored bacterial diversity patterns, whereas protozoan communities remained comparatively stable across experimental groups. These findings highlight the value of shotgun metagenomics for capturing multi-domain responses within the gut ecosystem and suggest that ecological perturbations associated with *S. mansoni* infection extend beyond bacterial community structure alone. Future studies should investigate the interactions among bacteria, bacteriophages, plasmids, and microbial eukaryotes to better understand their collective contribution to microbiome stability, genome evolution, and horizontal gene transfer.

Overall, this study demonstrates that IS elements are not only widespread in the gut microbiome but remarkably exhibit structured patterns of activity linked to host infection, microbial community composition, and genomic context. The preferential targeting of intergenic regions, combined with taxon-specific and condition-dependent variation in insertion activity, highlights the role of IS elements as key drivers of microbial genome plasticity in complex host-parasite-microbiota-associated ecosystems.

## Materials and Methods

### Sample origin, study design and ethics statement

The metagenomic dataset analysed in this study and obtained from intestinal contents of control/infected wild-type/human-microbiota-associated mice was generated in our previous study described in Cortés et al. ^15^. Briefly, microbiome-humanised and wild-type mice were either infected with *Schistosoma mansoni* or maintained as uninfected controls, following the experimental design reported previously ^15^. The metagenomic sequencing data are available from the European Nucleotide Archive (ENA) under study accession PRJEB40471.

### DNA extraction and sequencing

DNA extraction, library preparation, and shotgun metagenomic sequencing were performed as described in Cortés et al. ^15^. Briefly, total microbial DNA was extracted from faecal samples using the PowerSoil DNA Isolation Kit (QIAGEN) according to the manufacturer’s instructions. Sequencing libraries were prepared and sequenced on an Illumina NovaSeq platform, generating 150 bp paired-end reads. A full description of these procedures is provided in the original publication ^15^.

### Experimental workflow

The overall bioinformatic workflow used in this study is summarised in Fig. 7. Raw shotgun metagenomic sequencing reads were analysed using two complementary pipelines designed to characterise microbial community composition and insertion sequence (IS) activity. For taxonomic profiling, quality-filtered reads were classified using Kraken2, and species abundance estimates were refined using Bracken. The resulting taxonomic abundance tables were analysed in MicrobiomeAnalyst to assess microbial diversity and community structure across experimental groups.

For IS detection, metagenomic reads were assembled into contigs using MEGAHIT. The assembled contigs and processed reads were then analysed using the pseudoR pipeline to identify IS and estimate their abundance. Contigs containing IS were taxonomically classified using Kraken2 to determine the bacterial taxa associated with IS activity. In parallel, pseudoR and Prodigal were used to identify ORFs and determine whether IS occurred within coding (intragenic) or non-coding (intergenic) regions. Functional annotation of genes associated with IS-containing contigs was performed using eggNOG-mapper, allowing classification into functional categories and assessment of potential biological processes affected by IS mobilisation. Outputs from the taxonomic, insertional, and functional analyses were subsequently integrated to generate the final visualisations and comparative analyses presented in this study (Fig. S6).

### Bioinformatic processing of metagenomic data

Raw sequencing reads were processed using a quality control and preprocessing workflow adapted from ^15^. Trim Galore (v0.6.10) was used for quality and adapter trimming of whole-genome sequencing reads with default parameters. Paired reads in which at least one mate was shorter than 20 bp after trimming were discarded (https://github.com/FelixKrueger/TrimGalore). Quality filtered reads were mapped to the *Mus musculus* reference genome (GRCm38) using Bowtie2 ^42^ to remove host-derived sequences.

The resulting clean reads were classified using Kraken2 (v2.2.2) with the “--minimum-hit-groups” option, increasing the minimum number of hit groups from the default of two to three to improve classification accuracy against reference databases including bacterial, plasmid, fungal, plant, viral, and protozoan genomes ^43,44^. Species-level abundance in each sample was estimated using Bracken (v2.9) with default settings ^45^. Comparative analyses of gut microbiome composition in *S. mansoni* negative and positive HMA and WT mice were performed using MicrobiomeAnalyst ^46,47^. Following cumulative sum scaling (CSS) normalisation, differences in Shannon diversity and species richness between groups were assessed using unpaired t-tests or one-way or pairwise analysis of variance (ANOVA), as appropriate. Beta diversity was evaluated using PCoA based on Bray-Curtis dissimilarity matrices. Differences in community composition between groups were assessed using analysis of similarity (ANOSIM).

### Identification and analysis of insertion sequences

Duplicate removal and read decontamination were performed using tools from the BBTools software suite. PCR duplicate reads were removed using clumpify.sh with the parameter “subs=0”. Cleaned and deduplicated reads were assembled using MEGAHIT (v1.2.9) with the “meta large” preset (k max 127, k min 27, k step 10) ^48^. Only contigs longer than 1 kb were retained for downstream analyses. Insertion sequences were identified following the pseudoR workflow. Cleaned and deduplicated reads and assembled contigs were screened against the ISOSDB database using pseudoR and detected insertion sequences were catalogued for each sample according to the pipeline described in the original pseudoR publication ^21^. pseudoR identifies both the insertion allele, containing the IS element, and the corresponding non-insertion allele, representing the same genomic locus without the insertion. Relative IS depth was calculated as:

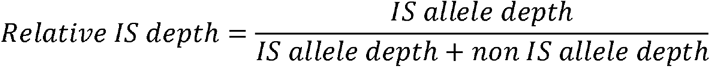

This metric estimates the proportion of reads supporting the insertion allele relative to all reads covering that locus.

### Estimating Insertion Sequence Enrichment in Intergenic vs Intragenic Loci

We quantified the preferential localisation of IS elements within intergenic and intragenic loci using an occurrence per million base pairs (OPM) normalised insertion rate. The OPM was calculated as:

Occurrence per million base pairs normalised insertion rate =

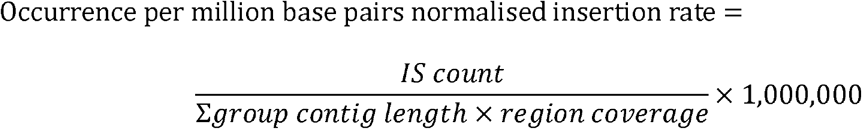

Where IS count represents the number of IS detected within a given genomic context (intergenic or intragenic), Σ group contig length is the total length of contigs assigned to the corresponding group, and region coverage denotes the proportion of the genome classified as intergenic (13%) or intragenic (87%) ^49^, respectively. This normalisation accounts for differences in assembly size and the relative genomic space available for insertions across groups.

### Functional and taxonomic classification of contigs and ORFs

Metagenomic sequences containing insertion sequences were functionally annotated using eggNOG mapper (v5) ^50^. Orthologous group assignments were cross-validated using the COG database. Open reading frames were classified as SusC, SusD, or TonB dependent if they contained PFAM domains matching the terms “SusC”, “SusD”, or “TonB”. Anti-MGE open reading frames were defined as those assigned to COG5340, COG2189, COG0286, COG0732, COG0827, or COG4217. Exopolysaccharide biogenesis and mobile element-related ORFs were assigned to COG functional categories “M” and “X”, respectively. Antimicrobial resistance genes were identified using the CARD Resistance Gene Identifier. Taxonomic classification of ORFs and contigs containing insertion sequences was performed using Kraken2 (v2.2.2) according to the pipeline described in the original pseudoR publication (Kirsch et al., 2024).

### Ethics

The life cycle of *S. mansoni* (NMRI strain) was maintained at the Wellcome Sanger Institute, Hinxton, UK, (WSI) by breeding and infecting susceptible intermediate hosts (*Biomphalaria glabrata* snails, NMRI strain) and definitive hosts (Female TO mice) as described ^20^. All experimental infections and regulated procedures from which samples and data were produced were approved by the Animal Welfare and Ethical Review Body (AWERB) of the WSI. All experiments were conducted under Home Office Project Licenses (Procedure Project License—PPL) No. P77E8A062 held by Gabriel Rinaldi, and No. P6D3B94CC held by Trevor D. Lawley. The AWERB is constituted as required by the UK Animals (Scientific Procedures) Act 1986 Amendment Regulations 2012.

## Acknowledgements

YHL was supported by a Computer Science Department Overseas PhD Scholarship (CSDOPS) and provision of high-performance computing resources used for data analysis at Aberystwyth University. GR was supported by UKRI Future Leaders Fellowships [MR/W013568/2].

## Author contributions

Conceptualisation: MTS, WA, GR

Formal analysis: YHL

Methodology and Investigation: YHL, MTS, WA, GR

Visualisation: YHL

Data curation: YHL

Supervision: MTS, WA, GR

Writing – original draft: YHL, MTS, WA, GR

Writing – review & editing: YHL, MTS, WA, GR, AC, CC

## Contributing interests

Authors declare that they have no competing interests.

## Data and materials availability

All data are available in the main text, accession numbers or the supplementary materials.

## Declaration of generative AI and AI-assisted technologies in the writing process

YHL declares that AI-assisted technology was used exclusively for proofreading and grammatical correction. All intellectual content, including study design, data analysis, and interpretation, was developed independently.

---

**Fig. S1:**
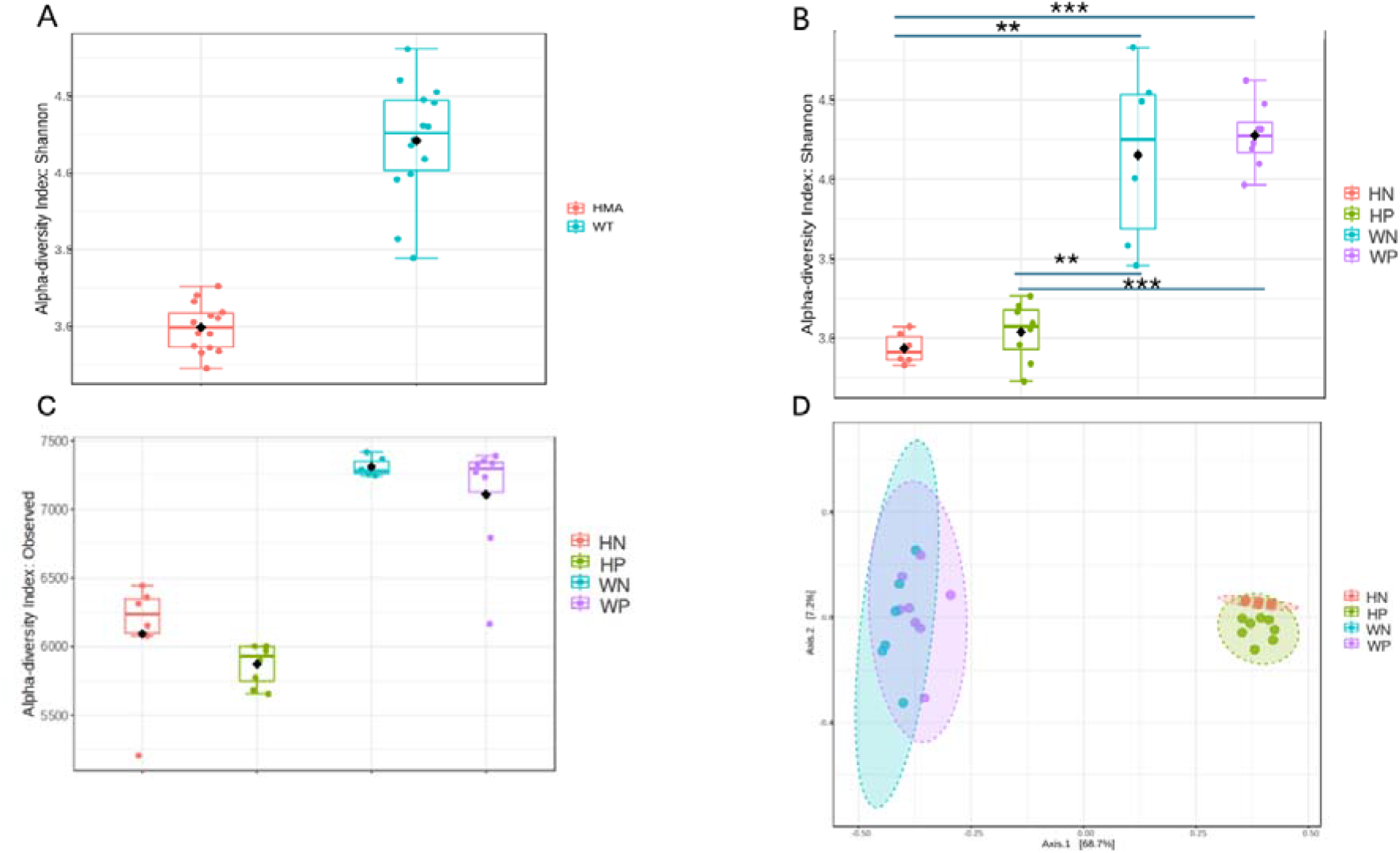
Bacterial compositional profiles of gut metagenomes in wild-type (WT) and human-microbiota-associated (HMA) mice. (A) Alpha diversity WT and HMA mice (Shannon index; t-test = −11.046, p = 2.86 × 10⁻⁹) (B) Alpha diversity of Schistosoma mansoni-infected and -uninfected WT and HMA mice (Shannon index; one-way ANOVA; F = 39.521, p = 1.92 × 10⁻⁹) Asterisks indicate significant differences between groups determined by pairwise ANOVA: *p<0.05; **p<0.001; ***p<0.0001 (C) Alpha diversity of Schistosoma mansoni-infected and -uninfected WT and HMA mice (Observed; one-way ANOVA; F = 34.939, p = 6.413 × 10⁻⁹) (D) Beta diversity of Schistosoma mansoni-infected and -uninfected WT and HMA mice (ANOSIM; R = 0.82813, p-value < 0.001).

**Fig. S2:**
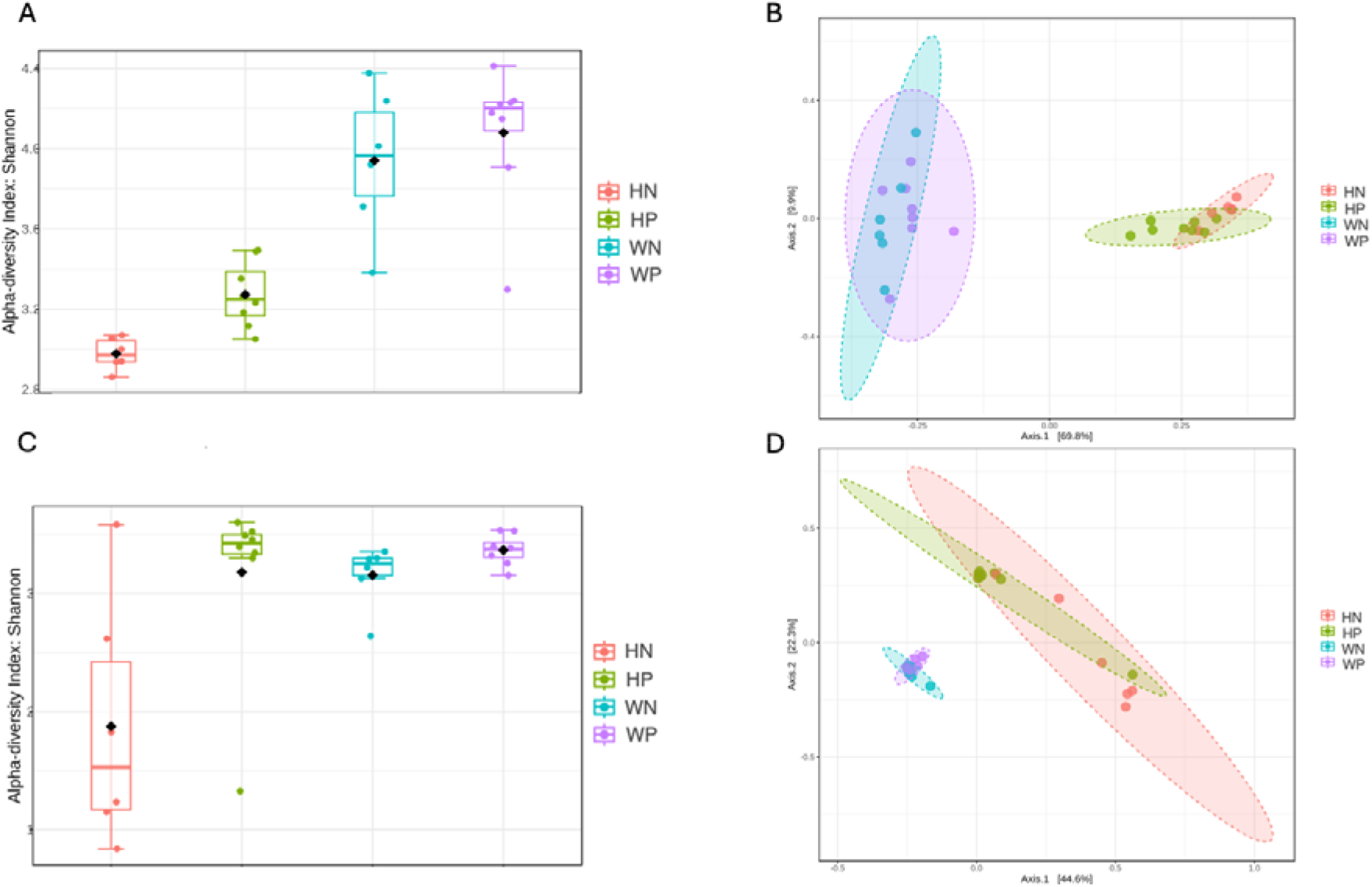
Plasmid and viral compositional profiles of gut metagenomes in wild-type (WT) and human-microbiota-associated (HMA) mice. (A) Plasmid alpha diversity of Schistosoma mansoni-infected and -uninfected WT and HMA mice (one-way ANOVA; F = 26.861, p = 7.63 × 10⁻⁸) (B) Plasmid beta diversity of Schistosoma mansoni-infected and -uninfected WT and HMA mice (ANOSIM; R = 0.78058, p-value < 0.001) (C) Viral alpha diversity of Schistosoma mansoni-infected and -uninfected WT and HMA mice (one-way ANOVA; F = 7.2507, p = 1.26 × 10⁻³) (D) Viral beta diversity of Schistosoma mansoni-infected and -uninfected WT and HMA mice (ANOSIM; R = 0.66645, p-value < 0.001)

**Fig. S3:**
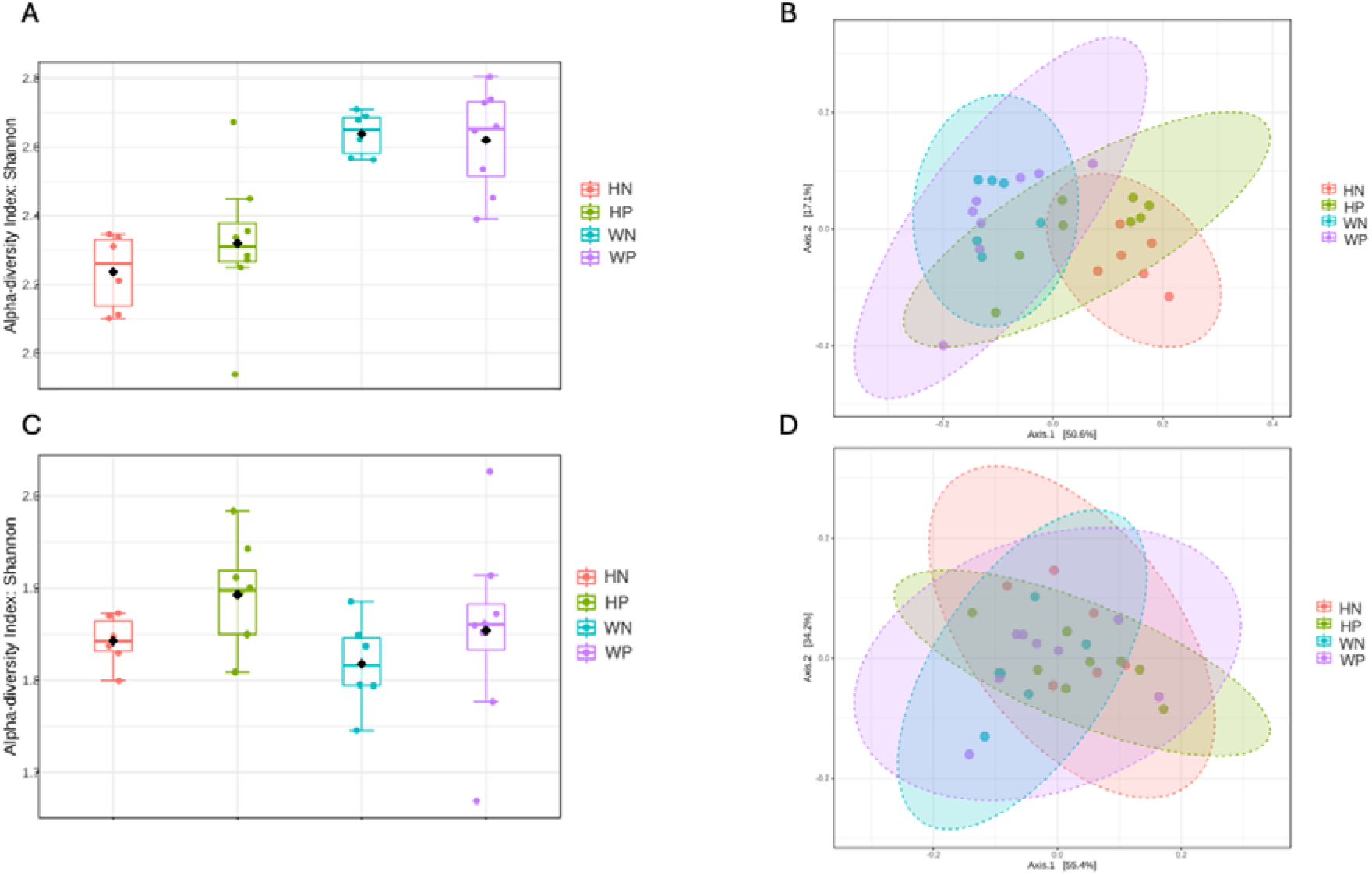
Fungal and protozoan compositional profiles of gut metagenomes in wild-type (WT) and human-microbiota-associated (HMA) mice. (A) Fungal alpha diversity of Schistosoma mansoni-infected and -uninfected WT and HMA mice (one-way ANOVA; F = 12.862, p = 3.27 × 10⁻⁵) (B) Fungal beta diversity of Schistosoma mansoni-infected and -uninfected WT and HMA mice (ANOSIM; R = 0.5454, p-value < 0.001) (C) Protozoan alpha diversity of Schistosoma mansoni-infected and -uninfected WT and HMA mice (one-way ANOVA; F = 1.4766, p = 0.246) (D) Protozoan beta diversity of Schistosoma mansoni-infected and -uninfected WT and HMA mice (ANOSIM; R = 0.17537, p-value < 0.01)

**Fig. S4:**
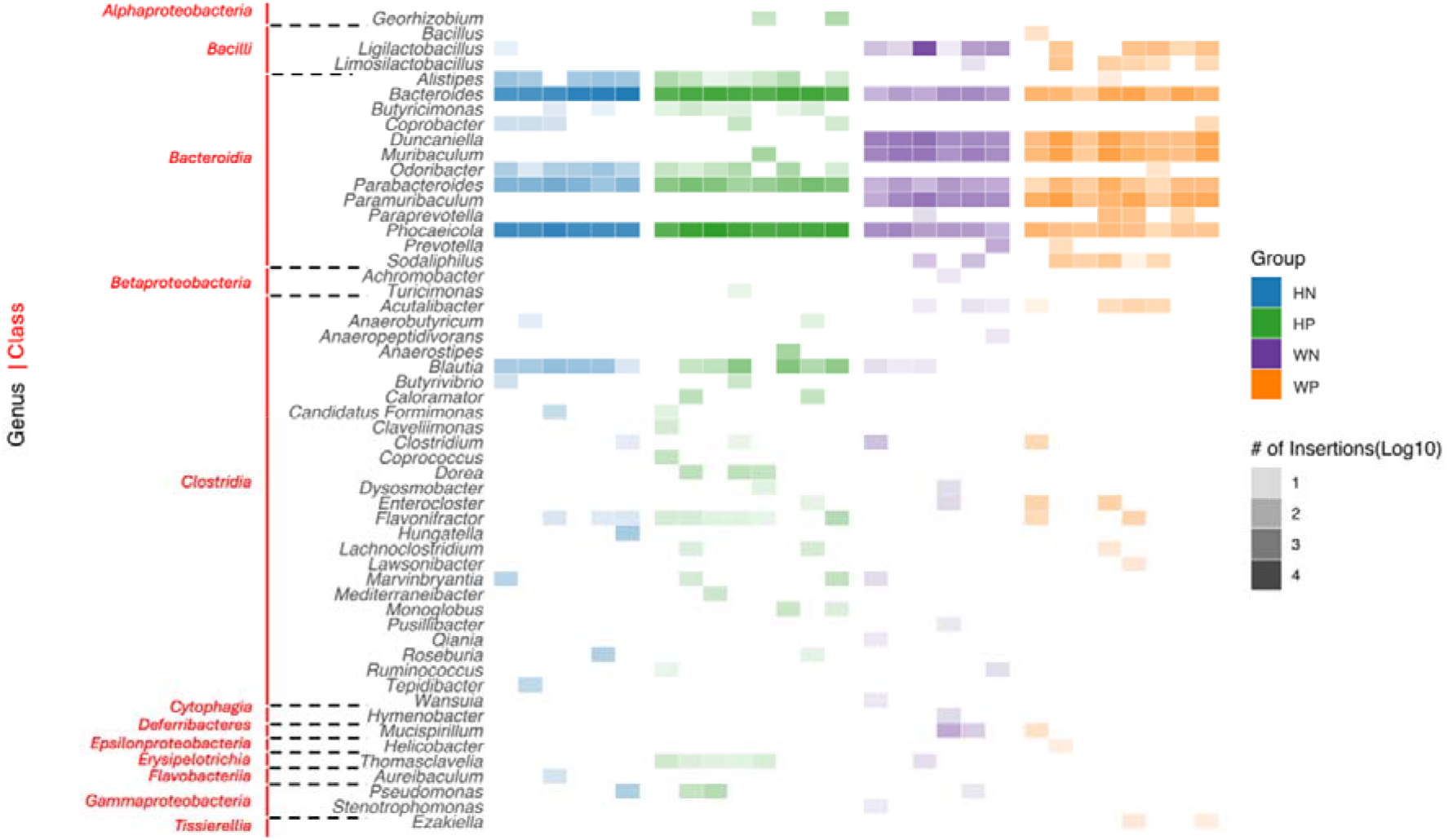
IS per bacterial class/genus. The intensity of each bar is proportional to the number of insertions

**Fig. S5:**
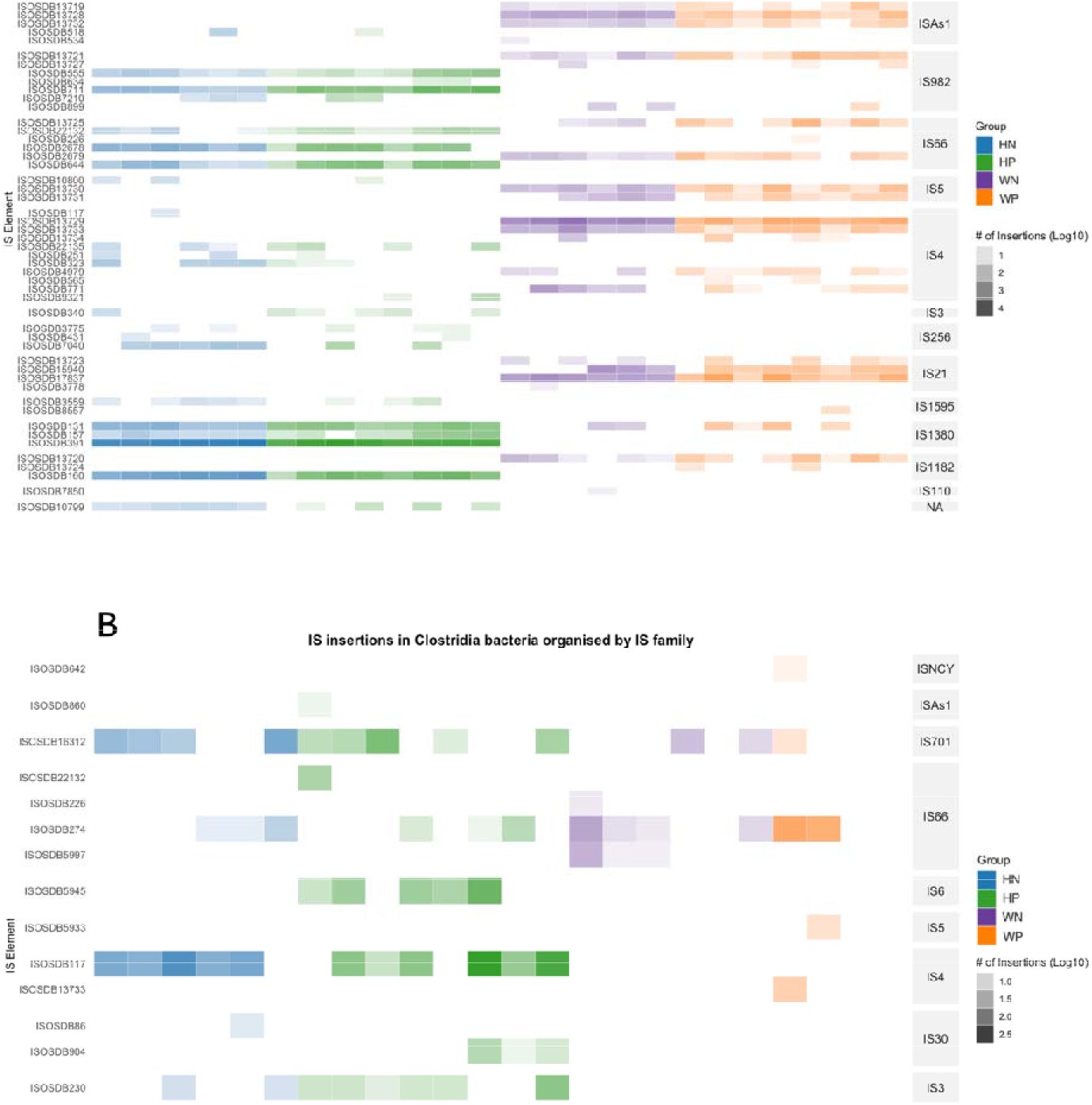
IS in (A) Bacteroidia and (B) Clostridia bacteria organised by IS family (right-hand side) with IS elements on the left. The intensity of each bar is proportional to the number of insertions

**Fig. S6:**
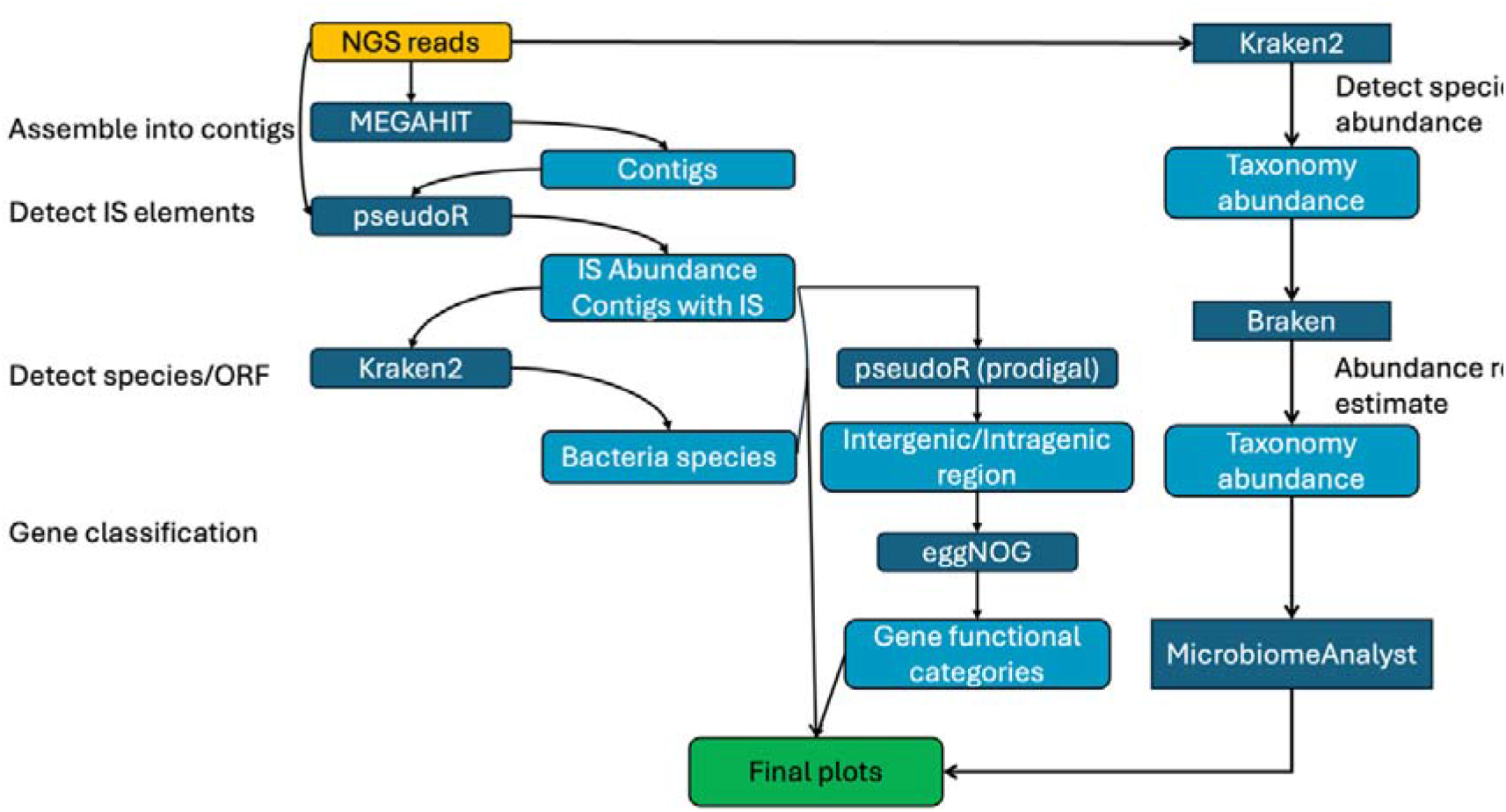
Overview of the bioinformatic workflow used to characterise microbial community composition and insertion sequence activity. Raw shotgun metagenomic sequencing reads were analysed using parallel taxonomic and IS detection pipelines. Taxonomic classification was performed using Kraken2 and Bracken, followed by downstream community analysis in MicrobiomeAnalyst. IS were identified using pseudoR following metagenome assembly with MEGAHIT. Contigs containing IS were taxonomically classified using Kraken2, while insertion locations and associated open reading frames were identified using pseudoR and Prodigal. Functional annotation was performed using eggNOG-mapper, and results from all analyses were integrated for downstream statistical analyses and visualisation.

